# Path integration error and adaptable search behaviors in a mantis shrimp

**DOI:** 10.1101/2020.03.04.977439

**Authors:** Rickesh N. Patel, Thomas W. Cronin

## Abstract

Mantis shrimp of the species *Neogonodactylus oerstedii* occupy small burrows in shallow waters throughout the Caribbean. These animals use path integration, a vector-based navigation strategy, to return to their homes while foraging. Here we report that path integration in *N. oerstedii* is prone to error accumulated during outward foraging paths and we describe the search behavior that *N. oerstedii* employs after it fails to locate its home following the route provided by its path integrator. This search behavior forms continuously expanding, non-oriented loops that are centered near the point of search initiation. The radius of this search is apparently scaled to the animal’s accumulated error during path integration, improving the effectiveness of the search. The search behaviors exhibited by *N. oerstedii* bear a striking resemblance to search behaviors in other animals, offering potential avenues for the comparative examination of search behaviors and how they are optimized in disparate taxa.

**Summary Statement:** Mantis shrimp use path integration, an error-prone navigational strategy, when travelling home. When path integration fails, mantis shrimp employ a stereotyped yet flexible search pattern to locate their homes.

## Introduction

Path integration is an efficient navigational strategy that many animals use to return to a specific location. During path integration, an animal monitors the turns it makes and distances it travels from a reference point using a biological compass and odometer. From this information, a home vector (the most direct path back to the reference point) is continuously updated, allowing the animal to return to its original location (Seyfarth et al., 1982; Muller and Wehner, 1988; Seguinot et al., 1993). Path integration is especially useful for central place foragers, animals which return to a home location between foraging bouts.

Due to small errors made in angular and odometric measurements during path integration, the home vector is prone to error accumulated over the course of an animal’s outward path (the path from the animal’s start location to the site of home vector initiation). Therefore, with a longer outward path, an increased error of the home vector is expected (Muller and Wehner, 1988; Cheung et al., 2007). To account for this error, some path-integrating animals initiate a stereotyped search behavior if they fail to reach their goal after following the path provided their home vector (Wehner and Srinivasan, 1981; Hoffmann, 1983; Durier and Rivault, 1999).

Many stomatopod crustaceans, more commonly known as mantis shrimp, are central place foragers that inhabit benthic marine environments. These animals occupy burrows in marine substrates, where they reside between foraging bouts (Dominguez and Reaka, 1988; Bach and Engle, 1989; Caldwell et al., 1989). Mantis shrimp of the species *Neogonodactylus oerstedii* employ path integration to efficiently navigate back to their burrows while foraging. However, the return paths guided by their home vectors often do not lead them directly to their burrows. When this happens, *N. oerstedii* initiate search paths to find their homes (Patel and Cronin, 2020). Here we investigate the source of home vector error in *N. oerstedii* and evaluate the means by which *N. oerstedii* copes with this error—its search pattern.

## Results and Discussion

### Path Integration in mantis shrimp is prone to accumulated error

In order to investigate the source of home vector error in *N. oerstedii*, individuals were placed in relatively featureless circular arenas in a glass-roofed greenhouse. Arenas had a sandy bottom and were filled with sea water. Vertical pipe burrows were buried in the sand so that they were hidden from view when experimental animals were away. Snail shells stuffed with small pieces of shrimp were placed at one of two fixed locations approximately 70 cm from the location of the burrow in the arena. Foraging paths to and from the location of the food were video recorded from above.

During these trials, animals would make tortuous paths away from the burrow until they located the food placed in the arena (the outward path). After animals located the food, they often executed a fairly direct homeward path (the home vector) before initiating a search behavior if their home vector did not lead them to the hidden burrow (Fig. 1A). Search behaviors were considered to be initiated when an animal turned more than 90° from the trajectory of its home vector. We defined the distance from the point of search behavior initiation to the location of the burrow the path integration error. We found that this path integration error correlates with outward path lengths during these trials (p = 0.017, r = 0.67, n = 12; Fig. 1B), suggesting that the error of path integration is an outcome of error accumulated over the course of the mantis shrimps’ outward paths.

**Figure 1.**
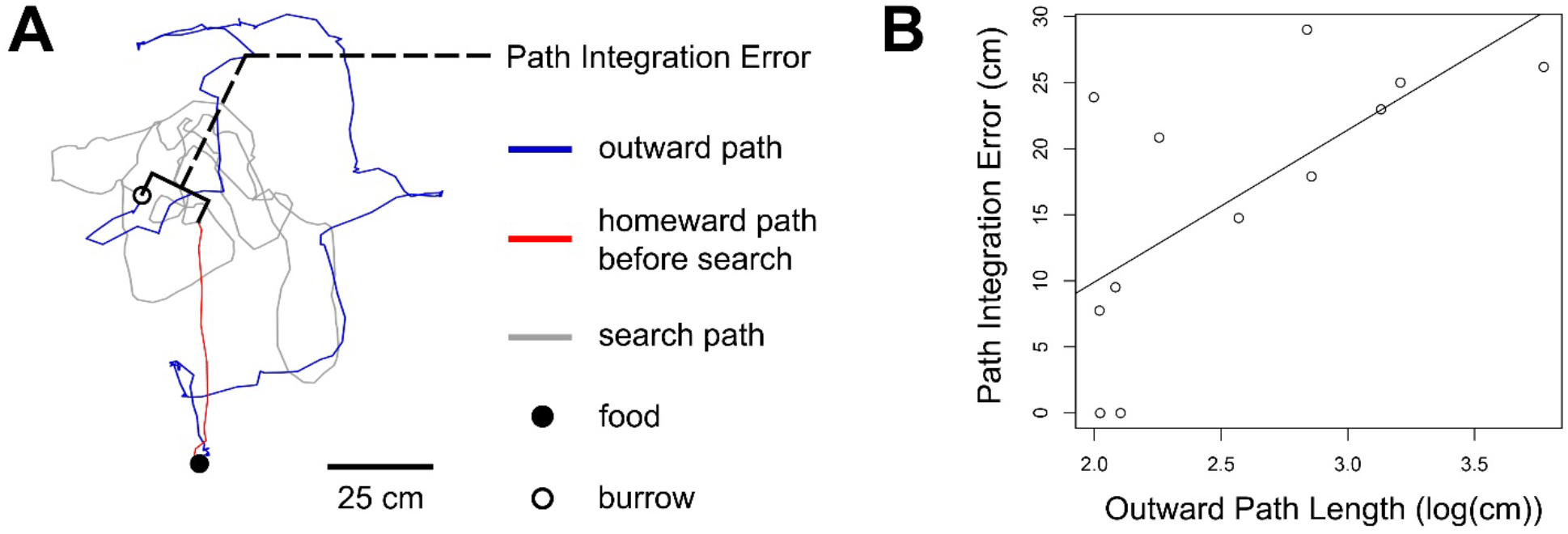
Error accumulated during outward forgaging paths leads to error in the home vector. **(A)** Example of a foraging path of *Neogonodactylus oerstedii*. The distance between the point search behaviors were initiated from the burrow location is the error of the animal’s path integrator. **(B)** Correlation between outward path lengths (log transformed) and the path integration error during trials in which the animals were not manipulated (P = 0.017, r = 0.67, n = 12).

### Search behaviors in N. oerstedii are stereotyped and flexible depending on error accumulated during path integration

Mantis shrimp execute stereotyped search behaviors when they have run through the paths provided by their home vectors without finding their burrows (Fig. 2, Supplemental Fig. 1, and Supplemental Video 1). These search behaviors are composed of loops that start and end near the location where the search is initiated (Fig. 2). We defined a loop in the search as a path that increased in distance from the point of search initiation before the animal turned and moved back toward the search initiation point. The loop was determined to be completed when an animal moved closest to the point of search initiation before once again moving away from the search initiation point or when an animal turned more than 90° from its trajectory back towards the search initiation point after returning halfway back to it, whichever occurred first (Fig. 2D).

**Figure 2.**
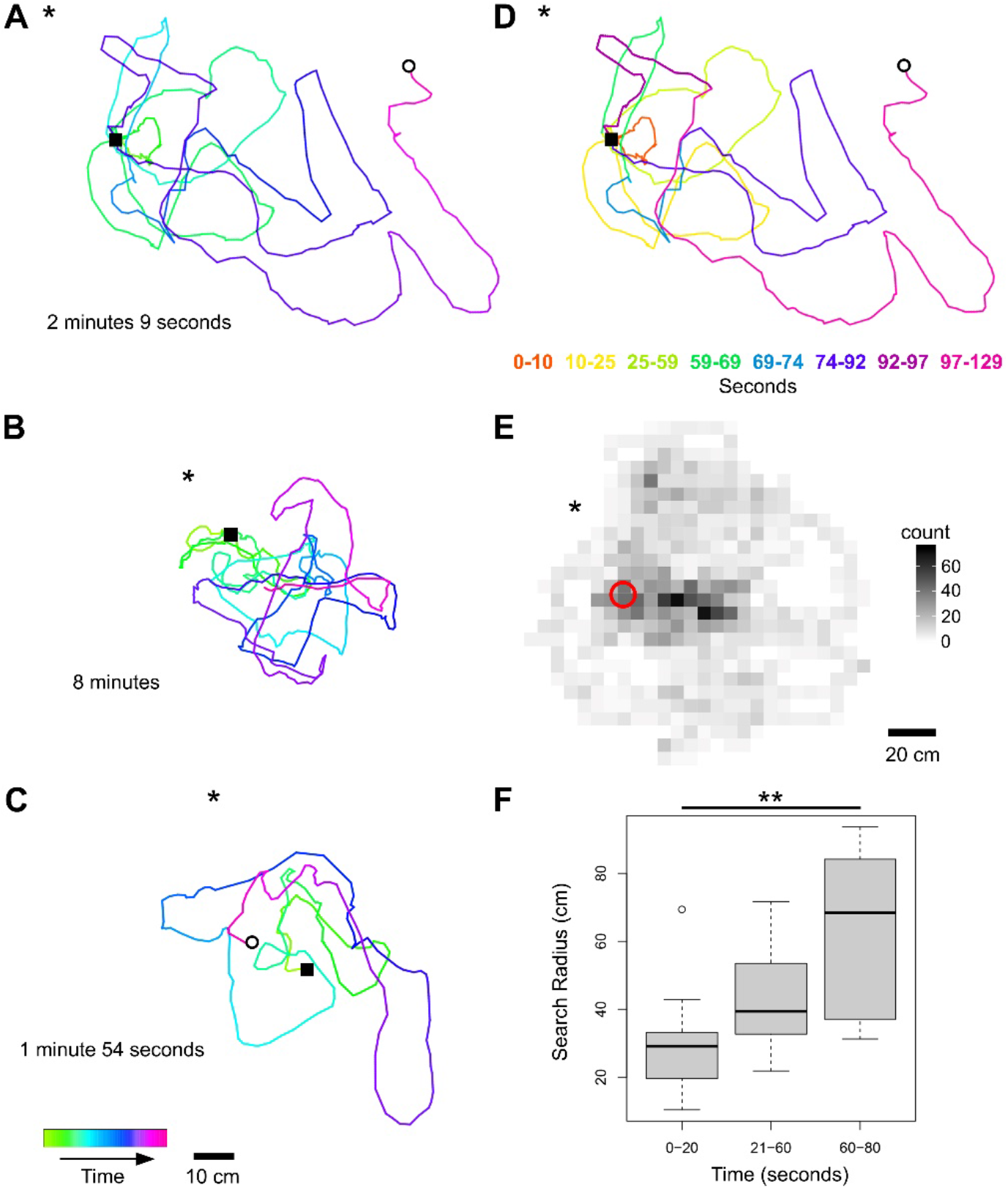
Search behaviors consist of a series of consecutive loops of increasing size which start near and return to a central location. Examples of search behaviors during trials when **(A and B)** animals were displaced before they initiated a home vector and **(C)** an animal was not manipulated. Empty circles represent the location of the burrow. Filled squares represent the location of search behavior initiation. Asterisks mark the location of the nearest edge of the arena. Lines are graded with time according to the key in the bottom left corner. **(A)** During this trial, the individual carried a food-filled shell during its homeward path and dropped it once it initiated its search behavior (marked by the filled square). This offered an opportunity to observe the strategy behind the search behavior, where consecutive continually-increasing concentric loops are made from the location of the initiation of the search behavior until the goal has been found. **(B)** This animal did not find its burrow after eight minutes of searching. The full search can be seen in Supplemental Fig. 1. **(D)** The same search as in (A) with search loops color-coded according to the key. **(E)** A heat-map of search behaviors complied from all trials in which animals were displaced in the arena (n=7). Shades of grey indicate counts of video frames in which animals moving more than one body-length per second were present at that location. Darker areas represent areas in the arenas where animals spent more time searching. The red circle marks the location of search behavior initiation and the asterisk marks the average nearest edge of the arena. Search behaviors are centered near the point of initiation. The observed deviation of the highest trafficked areas from the exact point of search behavior initiation might have been due to the initiation point’s proximity to the border of the arena. **(F)** The radii of search behaviors measured between 0-20 seconds, 21-60 seconds, and 61-180 seconds after search initiation. Search behaviors widen over time (ANOVA, P = 0.006, F = 6.374, 0-20 seconds: n = 11, 21-60 seconds: n = 10, 61-180 seconds: n = 7). Bars represent medians, boxes indicate lower and upper quartiles, whiskers show sample minima and maxima, and the dot represents and outlier (included in analyses). Asterisks indicate a significant difference in search radii between groups (P <0.01).

Successive search loops were not oriented between individuals (Supplemental Fig. 2). Further, all loops within single searches were not oriented in most individuals (Supplemental Fig. 3); however, searches in some individuals were biased away from the edge of the arena nearest to the location where the search was initiated (loops were only significantly oriented in two of eleven individuals: P = 0.03 and P = 0.025; Supplemental Fig. 1 and 3). These exceptions suggest that *N. oerstedii* can estimate its distance from a goal using local structures (here, the walls of the arena) and use these estimates to alter its searches in some cases.

**Figure 3.**
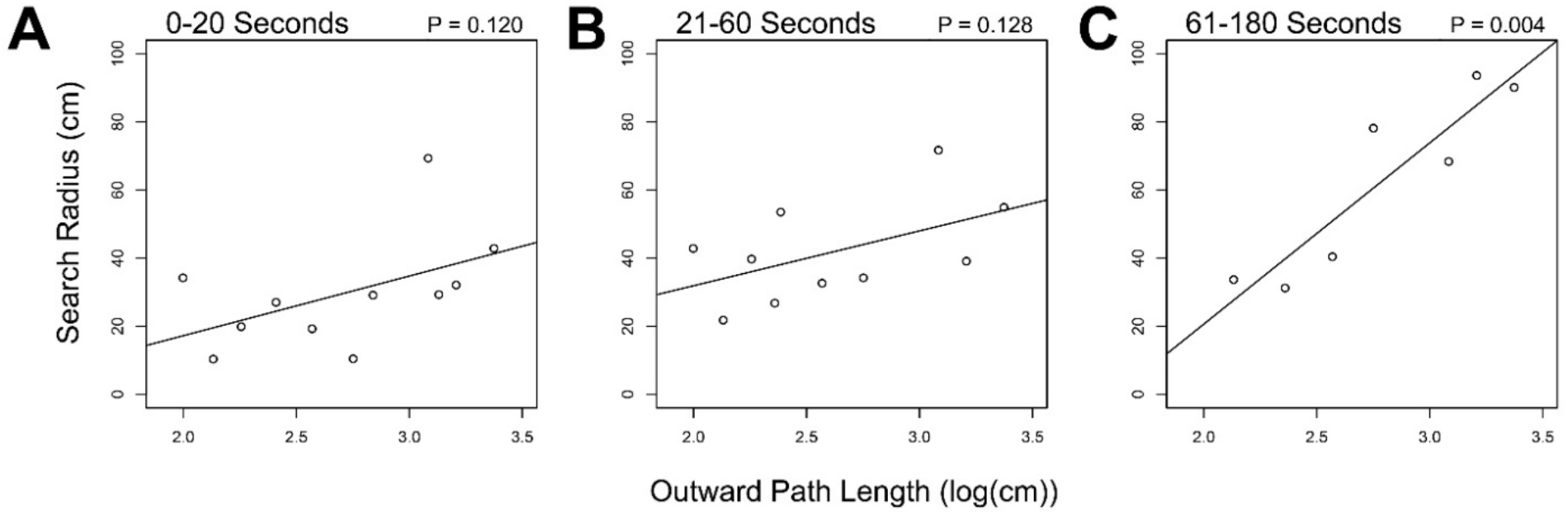
Search behavior radii are larger when outward path lengths are longer. Correlations of the radii of search behaviors and the lengths of outward paths (log transformed) measured between **(A)** 0-20 seconds, **(B)** 21-60 seconds, and **(C)** 61-180 seconds after initiation. For radii measured 61-180 seconds after search behavior initiation the relationship was significant (0-20 seconds: P = 0.12, r = 0.5, n = 11; 21-60 seconds: P = 0.128, r = 0.52, n = 10; 61-180 seconds: P = 0.004, r = 0.91, n = 7).

We also measured the radii of searches (the farthest distance of a search from the point of search initiation) within three time ranges (0-20 seconds, 21-60 seconds, and 61-180 seconds) and found that searches tend to increase in size over time (ANOVA, P = 0.006, F 6.374; Fig. 2F).

We found that the radii of searches were variable at similar time-points among searches, with many searches over three times as wide as other searches (Fig. 2F, Supplemental Fig. 1). We hypothesized that searches were wider when the error in a mantis shrimp’s path integrator was higher (perhaps because the animal’s confidence in its home vector’s accuracy was lower). In order to test this hypothesis, we compared the radii of search behaviors within three time ranges (0-20 seconds, 21-60 seconds, and 61-180 seconds) to the outward path lengths during those same trials. We found that outward path lengths and search radii were correlated and reached significance when the radii of searches were measured between 61-180 seconds after search initiation (Pearson’s correlation, P = 0.004, r = 0.52; Fig. 3). This result suggests that the sizes of search behaviors are modulated by the reliability of the path integrator.

## Conclusion

Path integration in *N. oerstedii* is inherently prone to error, which accumulates over the course of an animal’s outward path. Error due to distance estimates are expected to increase linearly with increasing outward path lengths. However, the magnitude of angular errors differ depending on the manner in which angular measurements are taken. If directional information is measured in relation to a stable compass heading or environmental feature, angular errors would be of a lesser magnitude than if angular information is measured from a previous rotational estimate, which would compound over the course of an animal’s journey. Some models of error accumulation during path integration suggest that due to this large accumulated rotational error, path integration over extended distances (such as those exhibited by bees and ants) would require the use of a stable compass reference during navigation (Cheung et al., 2007; Cheung and Vickerstaff, 2010; Heinze et al., 2018). This may be true for mantis shrimp as well; however, previous work suggests that mantis shrimp do rely on idiothetic orientation during path integration when celestial cues are obscured (Patel and Cronin, 2020). If mantis shrimp are indeed using idiothetic path integration when celestial information is unavailable, they would be relying on cumulative rotational estimates to measure their angular displacements under these conditions. Perhaps the typical limited foraging distances *N. oerstedii* exhibit in nature (usually not greater than a couple of meters) allow them to home using idiothetic path integration with reasonable accuracy.

To cope with the error in the home vector, *N. oerstedii* executes stereotyped search behaviors composed of a series of non-oriented loops (unless local features are detected) which increase in size over the course of the search. Even though these searches are stereotyped, their sizes are scaled: they become larger with increased outward path lengths, i.e. increased accumulated error during path integration. This flexible strategy improves the efficacy of the search.

In this study, some mantis shrimp searches were biased away from the edge of the arena nearest to where they initiated the search (Supplemental Fig. 1 and 3). This result suggests that mantis shrimp can judge the distance of a goal from nearby structures, which may act as landmarks. Similar search biases to local features have been observed in other animals. Desert ants alter the geometry of their searches for their nests depending on the apparent image size of the local landmark array on their retinas (Akesson and Wehner, 1997). Trained honeybees also have been demonstrated to use the apparent sizes of landmarks in their environment to focus their searches for a hidden food source (Cartwright and Collet, 1983). Landmark navigation is a reliable way for animals to correct for error accumulated during path integration and is often used by other animals in tandem with path integration to lead them to their targets (Etienne, 1992; Collett, 1996; Wehner, 2003). Mantis shrimps, many of which occupy structurally complex environments, may also use landmarks to assist their navigation.

The search behaviors of *N. oerstedii* closely resemble those executed by other animals, such as Catagylphid desert ants (Wehner and Srinivasan, 1981), cockroaches (Durier and Rivault, 1999), and desert isopods (Hoffmann, 1983). The searches of these animals are similarly composed of ever-expanding loops centered near the animal’s estimate of its shelter position and strikingly resemble the searches of mantis shrimp reported in this study. As in mantis shrimp, the radii of desert ant searches are also flexible (Merkle et al., 2006; Schultheiss and Cheng, 2011); however, in Cataglyphid ants, the search radii are scaled to the length of the home vector (Merkle et al., 2006), not to the length of the outward foraging path (Merkle and Wehner, 2010) such as we found in *N. oerstedii*. Due to the similarities of searches in these disparate groups of animals, the neural programs of these search behaviors may either be ancient homologs or remarkable convergences between insects and malacostracan crustaceans. Even if the underlying mechanisms of the searches these groups exhibit are homologous, differences in how these searches are manifested and elaborated are likely to be present. For example, the information used to scale search radii appears to differ between Cataglyphid ants and mantis shrimp.

## Materials and Methods

All data in this study were collected from experiments reported in Patel and Cronin (2020). Specifically, foraging behaviors from the “not manipulated” and “animal displaced” groups of trials enacted in the greenhouse on the UMBC campus in Patel and Cronin (2020) were used in the current study. All details of animal care, experimental apparatus, and experimental procedures can be found in Patel and Cronin (2020).

### Data and Statistical Analyses

Foraging paths from the burrow to find food and from food locations back to the burrow were video recorded from above. In order to differentiate homeward paths from continued arena exploration, paths from the food locations were considered to be homeward paths when they did not deviate more than 90° from their initial trajectories for at least one-third of the beeline distance (the length of the straightest path) from the food location to the burrow. From these homeward paths, search behaviors were determined to be initiated when an animal turned more than 90° from its initial trajectory.

Paths were traced at a sampling interval of 0.2 seconds using the MTrackJ plugin [15] in ImageJ v1.49 (Broken Symmetry Software), from which the output is given in Cartesian coordinates. From these data, the lengths of outbound, homebound, and search paths were calculated. The distance of the point of search behavior initiation to the burrow (the path integration error) was also measured using the MTrackJ plugin. Path integration error analyses were only measured from trials when animals were not manipulated.

Search behaviors lasting over ten seconds with at least one completed loop were analyzed from all trials when animals were either not manipulated and or were displaced to a new location in the arena (n = 11). We defined a loop in the search as a path that increased in distance from the point of search initiation before the animal turned and moved back toward the search initiation point. The loop was determined to be completed when an animal moved closest to the point of search initiation before once again moving away from the search initiation point or when an animal turned more than 90° from its trajectory back towards the search initiation point after returning halfway back to it, whichever occurred first.

The radii of search behaviors were measured as the farthest distance of a search from the original point of search initiation (i.e. the end point of the home vector) using ImageJ. The radii of searches were measured over three time ranges after search initiation: 0-20 seconds, 21-60 seconds, and 61-180 seconds. Additionally, orientations of search loops when loops were at the farthest distance from the search initiation point were recorded using ImageJ. For these measurements, searches were oriented so the axis of each home vector preceding the searches was at 0 degrees.

All statistical analyses were run on R (v3.3.1, R Core Development Team 2016) with the “CircStats”, “circular”, “plotrix”, “Hmisc”, and “boot” plugins.

Rayleigh tests of uniformity were used to determine if loops within individual searches had a directional bias and if successive search loops between individuals contained some pattern of orientation (Batschelet, 1981). All reported mean values for orientation data are circular means. All circular 95% confidence intervals were calculated by bootstrapping with replacement over 1000 iterations.

An Analysis of Variance Test (ANOVA) was used to determine if the radii of searches differed between the time intervals measured. A Tukey Honest Significant Difference post-hoc analysis was used to determine significant differences between groups.

Parametric Pearson’s correlation tests were used for all correlative analyses when applicable. For distributions that did not meet the assumptions of a Pearson’s correlation test, a non-parametric Spearman’s rank-order correlation test was used instead.

## Author Contributions

R.N.P. designed and conducted all research, analyzed all data, and prepared the manuscript. T.W.C. provided guidance and research support.

## Data and Materials Availability

The data that support the findings of this study are available from the corresponding author upon reasonable request. Correspondence and requests for materials should be addressed to R.N.P.

## Supplemental Figures

**Supplemental Figure 1.**
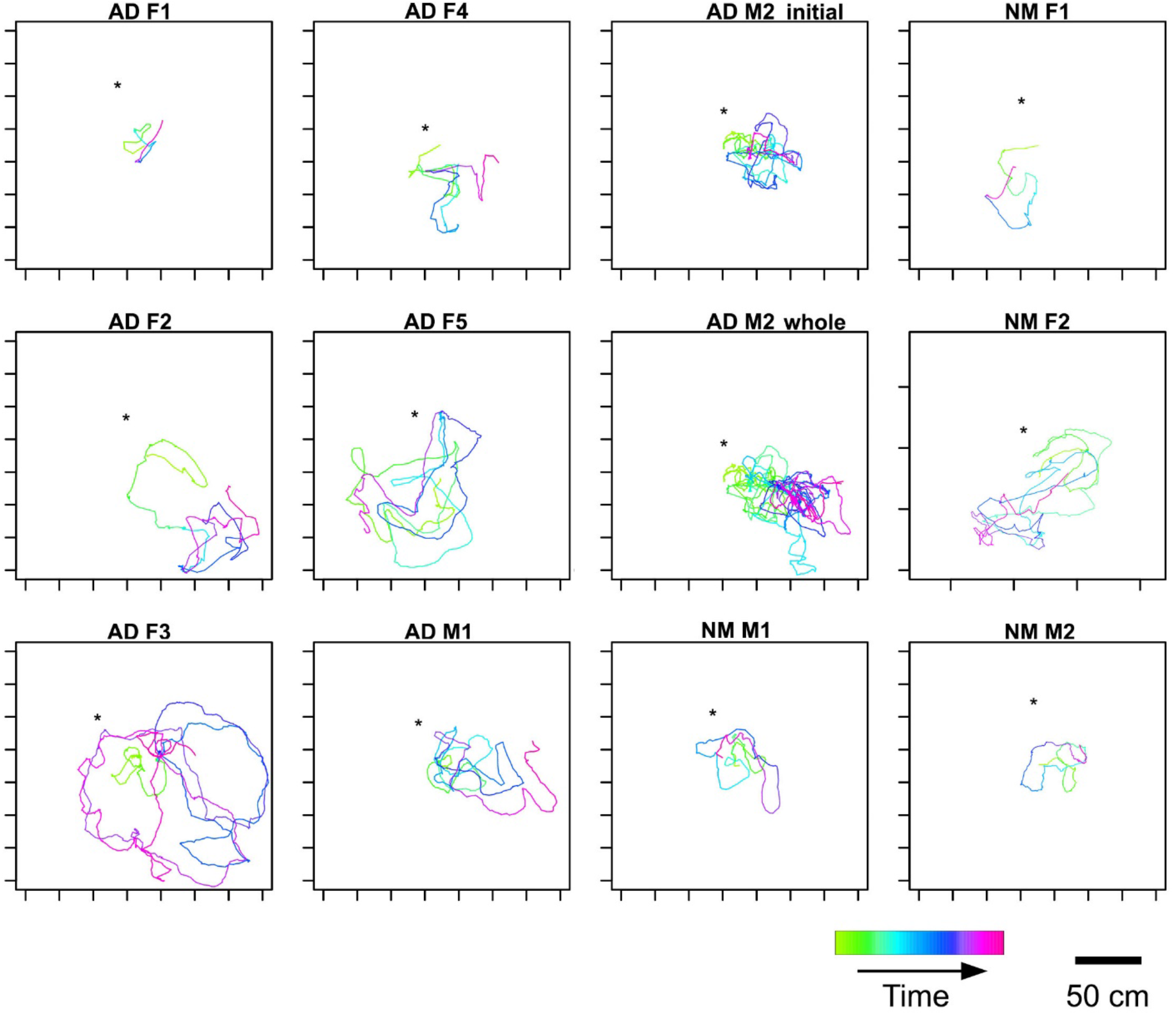
Path tracings of all searches. Asterisks mark the location of the nearest edge of the arena. The home vectors preceding the searches are oriented vertically. The search for AD M2 is plotted as both, the first eight minutes of the search (initial) and the whole search (whole).

**Supplemental Figure 2.**
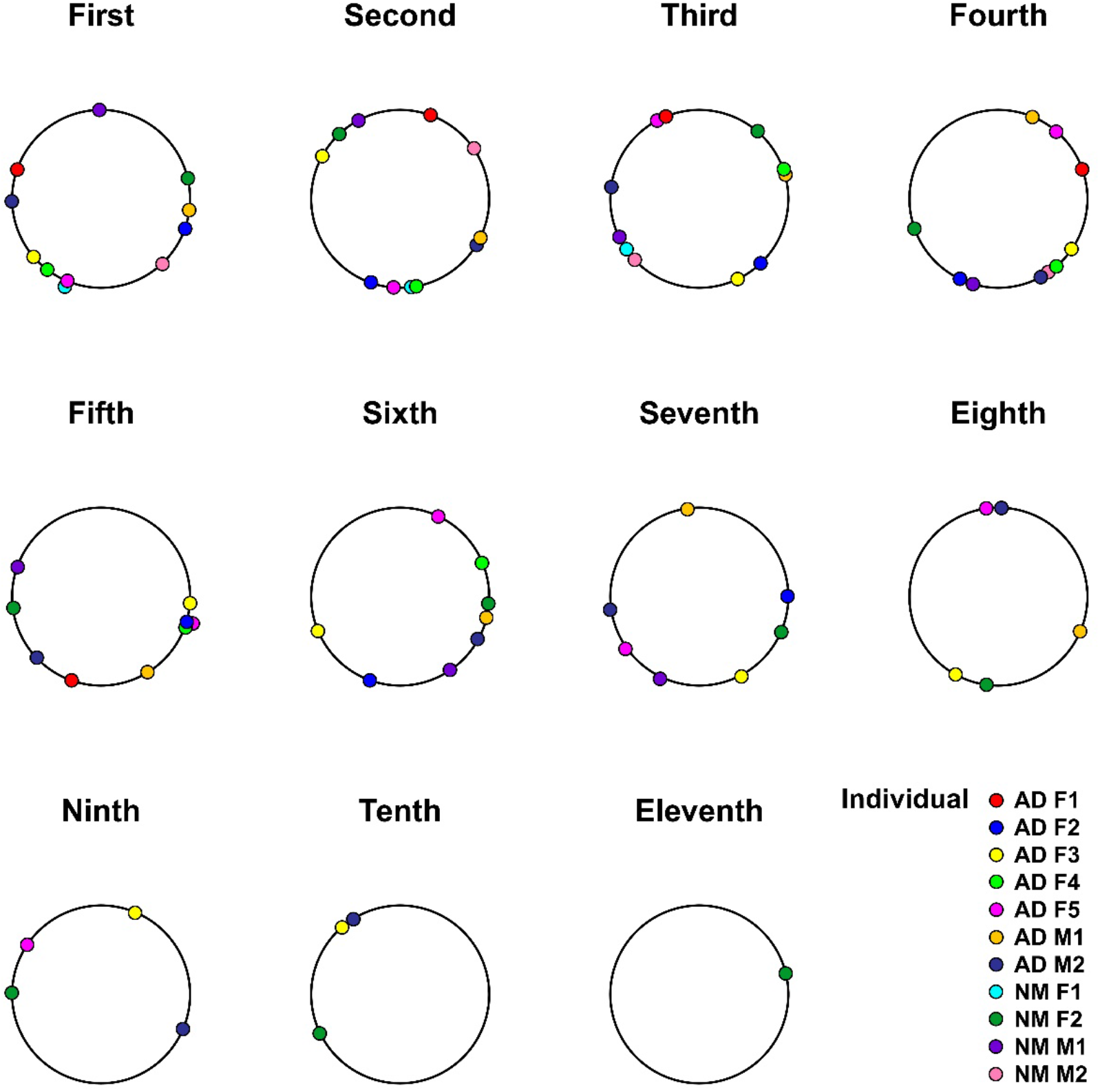
Search loop orientations per loop number. Successive search loops were not significantly oriented between individuals. The home vectors preceding the searches are oriented to the top of the plot (towards 0 degrees).

**Supplemental Figure 3.**
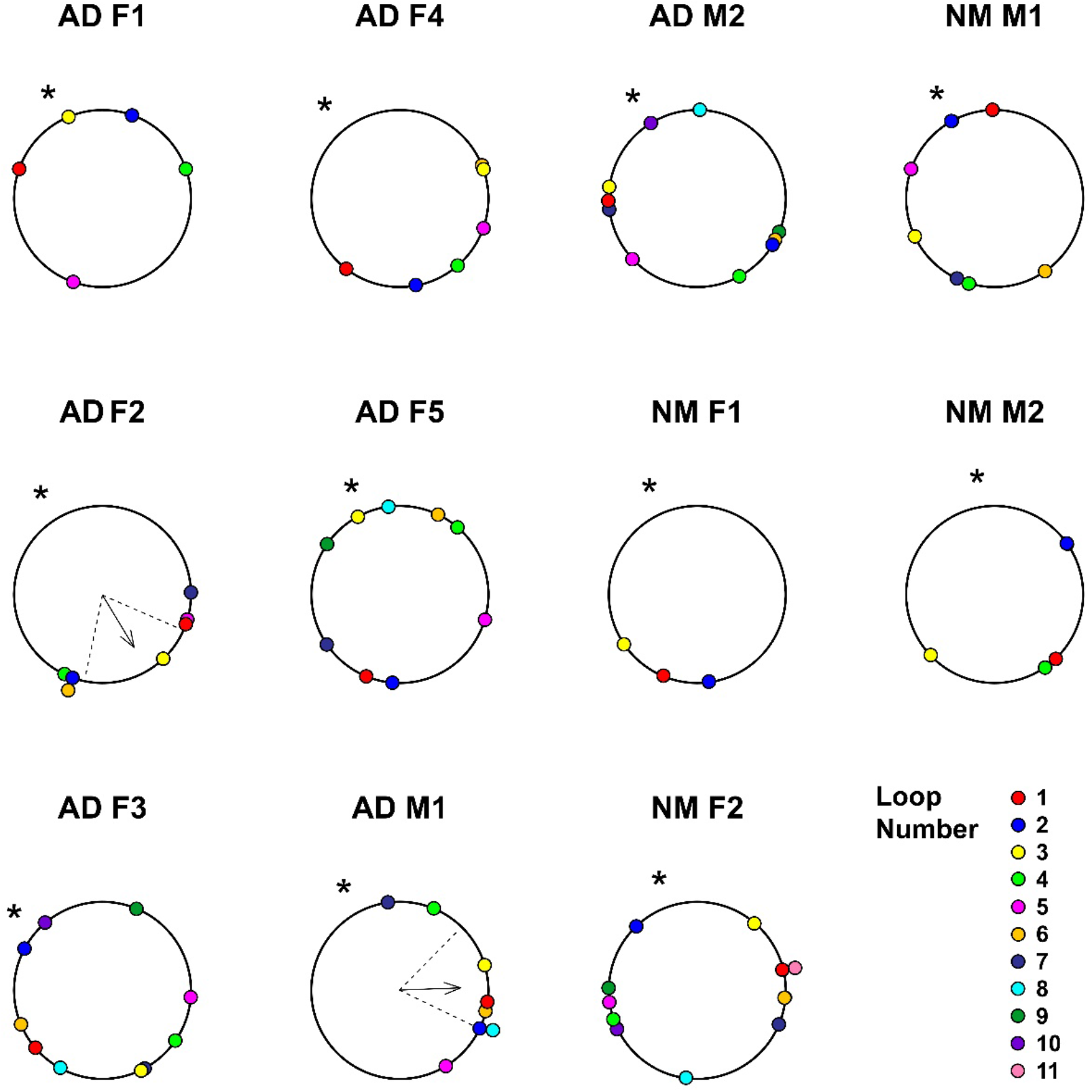
Search loop orientations per individual. Asterisks mark the direction of the nearest edge of the arena. The home vectors preceding the searches are oriented to the top of the plot (towards 0 degrees). Arrows in plots represent mean vectors, where arrow angles represent vector angles and arrow lengths represents the strength of orientation (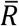). Dashed lines represent 95% confidence intervals. Means and 95% confidence intervals were only included in plots with significant orientations (AD F2: P = 0.03, 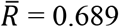 and AD M1: P = 0.025, 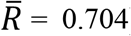). Loops appear to be biased away from nearest edge of the arena in these individuals.

**Video S1. Search behavior of *Neogonodactylus oerstedii* following a home vector.** Home vector path is in red and search path is in grey. This animal was displaced preceding the home vector and search seen in the video. The search path tracing can be reviewed in Figure 2B. Filmed at 30 frames per second. Replay speed is indicated in the bottom-right corner of the video.

